# Exploration of Antibacterial Activities of *Berberis royleana* Fractions Extracts

**DOI:** 10.1101/2021.06.14.448462

**Authors:** Muhammad Rafique, Muhammad Salman

**Affiliations:** Faculty of Life Sciences Department of Microbiology and Biotechnology, Abasyn University Peshawar

## Abstract

**Objectives:** Prepare various solvent extracts of *Berberis royleana* (areal part of plant) to determine the *in vitro* antibacterial potential of methanolic, ethyl acetate, chloroform, *n*-hexane and water extracts of *B. royleana* against various bacterial isolates and Compare the efficacy of outstanding antimicrobial extracts of *B. royleana* with commonly used antibiotics. *Berberis* species are medicinally important plants, produce various metabolites and used as treatment for multiple complications. *Berberis royleana* is a rare specie belongs to genus *Berberis*.

**Methods:** In the current study the areal parts of the plant were isolated to explore antibacterial activities. Antibacterial activities were done using standard procedures. The antibacterial activities of different fractions were tested by 100 μg methanolic, ethyl acetate, chloroform, *n*-hexane and water fractions of *B. royleana* against bacterial isolates *Escherichia coli, Staphylococcus aureus, Klebsiella pneumoniae, Pseudomonas aeruginosa, Salmonella* Typhi and *Proteus spp*. The ciprofloxacin (5μg) was used as a positive control and DMSO as a negative control.

**Results:** All fractions showed zone of inhibition against the growth of tested bacterial isolates. Methanolic fractions have maximum ZI against *S. aureus* and *K. pneumoniae* (25.7±1.5 mm), *S. aureus* (23±2.7), *Salmonella* Typhi (25±1), water fraction have *Klebsiella pneumoniae* (24.4 ±2.5), *Salmonella* Typhi (23 ±1 mm), *S. aureus* (21±2.8 mm) and the *n*-hexane fraction exhibits ZI against *K. pneumoniae* (24.7±1.5), *Salmonella* Typhi (24±2) *S. aureus*, ethyl acetate maximum zone against *E. coli* (16.6±3.1) and chloroform fraction showed maximum ZI against *S.* Typhi (21.6±3.3).

**Conclusion:** From current report it may be concluded that *B. royleana* extracts have medicinally effective potentials against drugs resistant bacteria.

## I. INTRODUCTION

Multiple antimicrobial agents show resistance in pathogenic bacteria which become an important public health problems. For treatment of infection there are rare, or even sometimes no effective antimicrobial agents available. This justifies the exploration of alternative forms for the treatment of infections, such as novel compounds with bactericidal properties from natural sources like plants [1] [2]. The crude extracts of plant segments and phytochemicals of identified antimicrobial properties have importance in therapeutic cure [3]. The infectious microorganism have shown resistance to several antimicrobial agents due to environmental advancement and for this purpose new antimicrobial drugs are needed from time to time treat disease [4].

### 1.2 Medical uses of medicinal plants

The practice of traditional medicine is widespread in Pakistan, Japan, China, India, Sri Lanka and Thailand [5] [6] [7]. For treatment and curing of various human diseases and infections used variety of plant and different parts of plant are used in traditional medicine which is one of the oldest systems of world [8] [9] [10].

#### 1.2.1 Medicinal Plant against Bacterial Infection

The bacterial resistance is rising a problem and the perspective of the use of antimicrobial drugs in the future is extremely uncertain. Many pathogens cause diseases shows resistant to antibiotics as a result cause illness, death and an even huge economical loss [11]. Increasing levels of antibiotic resistance in pathogenic bacteria has increased the demand to search novel antibiotics for diseases control. In mid-21st century due to the drug resistant there are millions of deaths occurred and also loss of trillions of dollars in recent reports [1]. The rise of antimicrobial resistance show in bacteria. The Gram-positive bacteria mostly cocci like coagulase-negative *Staphylococcus aureus*, *Enterococcus* and *staphylococci* species are crucial microorganisms in the hospital atmosphere [2] [12]. In the Enterobacteriaceae family Gram-negative bacteria are significant causes of urinary tract infections (UTIs), pneumoniae, invasive and various peritoneal infections. *K. pneumoniae* is second most causative pathogens which causing invasive infection after the *Escherichia coli*. A major resistance problem in Enterobacteriaceae infections are production of extended-spectrum b-lactamases (ESBLs) (David *et al*., 2006; [13].

##### 1.2.1.1 Escherichia coli

*Escherichia coli* (*E. coli)* is involved in both commensals and pathogenic strains of the gastrointestinal tracts of vertebrates which is causative pathogen for multiple extra and intra intestinal infections [14].

##### 1.2.1.2 Pseudomonas aeruginosa

*Pseudomonas aeruginosa* (*P. aeruginosa*) is a cosmopolitan Gram-negative aerobic bacillus, mainly nosocomial pathogen [15]. *P. aeruginosa* most common pathogen isolated from surgical site infection, invasive infection, nosocomial pneumonia, urinary tract infection (UTIs) [16]. *P. aeruginosa* caused diverse diseases which are existence of multiple pathogenic mechanisms must be expected [17]. In United States the *P. aeruginosa* bacteria are common cause of morbidity and also cause mortality in hospitalized patients [18].

##### 1.2.1.3 Klebsiella pneumoniae

*Klebsiella pneumoniae* (*K. pneumoniae*) is a Gram-negative bacteria, Cylindrical in shape and rod like. It is of about 2 microns in length and 0.5 microns in diameter and widely distributed in nature, found abundantly in soil and water [19]. *K. pneumoniae* is among the most common pathogenic bacteria and hospital-acquired pathogen which causing urinary tract infections (UTIs), pneumonia and peritoneal infection in hospitalized immune compromised patients with severe underlying diseases. It was also a possible community acquired pathogen and second only to *E. coli* in gram negative bacteria [20] [13]. Multiple antimicrobial agents are increasingly resistant to *K. pneumoniae* isolate which are including Quinolones and production of extended-spectrum Cephalosporins (ESC) extended-spectrum beta-lactamases (ESBL) such as Ceftazidime [21].

##### 1.2.1.4 Salmonella

In several countries resistant reported in *salmonella* species to expanded-spectrum Cephalosporins. *Salmonella* infections that occur in United States each year about 1.4 million and most are in the elderly, children and approximately 600 are fatal [22] [23].

##### 1.2.1.5 Proteus

*Proteus* species belong to family Enterobacteriaceae a bacillus which is Gram negative bacteria and also serious cause of infection in human [24]. Proteus species among those pathogens which can cause both community and hospital acquired infections. Because different mode of transmission it can causes various type of infection in human. Source of infection contaminated water, soil, food, equipment, catheters, intra venous lines, the hands of patients and health staff [25].

##### 1.2.1.6 Staphylococcus aureus

*Staphylococcus aureus* (*S. aureus*) is a commensal organism in human being which carried in the nares of 30% of healthy adults. *S. aureus* can colonize and cause a variety of supporative diseases in man including skin, blood, wound sites, bone infections and vascular catheters as well as the nasopharynx in the hospitalized patients [26] [27]. *S. aureus* is a hospital acquired diseases, serious community and a major problem of Public health [28].

### 1.3 Berberis royleana

*Berberis royleana* (*B. royleana*) is rare species amongst the members of *Berberis* (Berberidaceae). This specie is still imperfectly known and the flower is not specified. It differs from other species of the genus by its smaller leaves, narrow fruits, inflorescence and pedicels. *B. royleana* is a deciduous plant with height up to 3 m. The roots are thick and broaden easily. The stem has red-brown color and has spines. Leaves are usually 7-15 mm long, 6-12 mm broad. The ripe fruit is ovoid with red color and about 1 cm in length. The fruits developed in clusters form are bitter to taste. Berries are somewhat black, pruinose grey, oblong, 8 mm long and 3.5 mm broad immature [29].

#### 1.4.1 Medicinal uses of Genus *Berberis*

*Berberis* have been supportive within cure of malignancy, diabetes, jaundice, spleenomegaly, AIDS, arthritis, cardiac diseases, eye infections, hypertension, infectious diseases, cholera, diarrhea, wounds, intestinal colic, dysentery, amoebiasis, eye troubles, leprosy, antimalaric, anti-inflammatory, fever, inflammation (swelling), gastric pain, UTI infection, throat infection, gums infections [30][11][31] [32].

#### 1.4.5 Properties of *Berberis*

The roots of *Berberis* species are working as an anti-periodic, diaphoretic and antipyretic and its action was supposed to be as powerful as quinine. The shoot is used as a tonic, anti-periodic, cardiovascular, hepato-protective, antimicrobial and anti-cancerous activities [33].

## II. METERIAL AND METHOD

### 3.1 Study area

This study were carried out in Microbiology and Biotechnological Department Abasyn University Peshawar from February 2018 to July 2018.

### 3.2 Collection and identification

The Disease free *Berberis royleana* plant was collected from Azad Kashmir, Pakistan. The plant specimen was identified by Dr. Taj Ur Rahman, Department of chemistry Mohi-Ud-Din Islamic University Azad Jammu & Kashmir.

### 3.3 Sample preparation

The areal part of *B. royleana* plant was washed with distilled water. The *B. royleana* were air-dried at room temperature under shade. The plant was collected and powder was made by using electric grinder. The powder samples of plant were then packed in clear polythene pouches, sealed and stored at room temperature.

### 3.4 Preparation of crude extract of *B. royleana*

One kilograms powder of *B. royleana* was dissolved in 1000 ml methanol and kept at room temperature for 7 days. The mixture was intervally shaked to get maximum extract. The methanol soluble components was filtered using Whattman filter paper stored in container and the process was repeated for another 7 days. The extracts were collected from both containers and combined. The extract solution was dried using a vacuum pump with the rotary evaporator with reduced pressure at a temperature of 45 °C. The remaining methanol are evaporated at 45 °C in water bath. The semi-solid crude extract was obtained and kept in sterile bottles at 4°C until use.

### 3.5 Fractionation

About 55 grams of crude extract of *B. royleana* suspended in 200 ml distilled water and shaked with help of electromagnetic shaker. Methanolic crude extract was dissolved completely in distilled water and pour in separatory funnel. For further fractionation different solvents was added into the mixture. I.e. *n*-hexane, ethyl acetate and chloroform [34] [35].

### 3.6 Stock concentration of extracts

All *B. royleana* fractions (methanol, water, *n*-hexane, ethyl acetate and chloroform) are were dissolved in 10 mg/1ml dimethyl sulfoxide (DMSO) solution for further use.

### 3.7 Control

Ciprofloxacin (5 μg) antibiotic was used as a positive control for both Gram positive and Gram negative bacterial isolates. Dimethyl sulfoxide (DMSO) solution was used as a negative control for antibacterial.

### 3.8 Isolation of tested microorganisms

#### 3.8.1 Bacterial isolates

The different bacterial isolates were used in the study such as *Staphylococcus aureus, Escherichia coli, Salmonella* Typhi*, Pseudomonas aeruginosa, Klebsiella pneumoniae* and *Proteus spp.* All the bacterial isolates were clinically obtained from Lady Reading Hospital (LRH) Peshawar, Hayatabad Medical Complex MTI (HMC) and MMC General Hospital Peshawar Khyber Pakhtunkhwa, Pakistan. Bacterial isolates were sub cultured on nutrient agar medium and incubated at 37 oC for 24 hours. The Broth of each bacterium was prepared from developed (24 hours) culture of bacteria on nutrient agar.

### 3.9 Antibacterial activity

#### 3.9.1 Preparation of nutrient agar and sampling

Sterilization of glass wares was done through steam sterilization using autoclave. Culture media was prepared in conical flask having distilled water using standard protocol. Nutrient agar (28 g) was dissolve distilled water made up volume up to 1000 ml followed by autoclaving at 121 °C for 15 min. The media was cooled upto 45 °C and poured into sterile Petri dishes and allowed to solidify at room temperature.

#### 3.9.2 Antibacterial activity of *Berberis royleana* extract

Antibacterial activity of the crude extracts was checked using agar well diffusion method by Aliakbarlu *et al*., 2014 [36]. Mueller-Hinton agar (38 g) was dissolve in distilled water made up volume 1000 ml and autoclaved. The autoclaved Mueller-Hinton agar was poured in Petri dishes and allowed to solidify for 30 minutes. Mueller-Hinton agar plates were kept in Laminar Flow Hood and bacterial culture (*Staphylococcus aureus, Escherichia coli, Salmonella* Typhi*, Pseudomonas aeruginosa, Klebsiella pneumoniae* and *Proteus spp*) was inoculated using sterile cotton swabs to achieve uniform lawn of growth. Wells of 6 mm in diameter and about 20 mm apart were punctured in the culture media using sterile cork borers to make three to five uniform wells in each Petri dish. A drop of molten nutrient agar was used to seal the base of each well. The 100 μg of each fraction of *B. royleana* plant extract (methanol, water, *n*-hexane, ethyl acetate, chloroform) was introduced through micropipette aseptically into specifically marked wells in the agar plates and the antibiotic (Ciprofloxacin 5μg) was used as positive control for specific bacteria and dimethyl sulfoxide (DMSO) as a negative control. The plates were kept undisturbed for 45 minutes to make the free diffusion of the extract. The antibacterial activities were observed after 24 hours of incubation at 37 °C. The zone of inhibition by fraction extracts of *B. royleana*, positive and negative controls were measured (mm) and snapshots of inhibitory zone of all bacterial isolates were saved.

## IV. RESULTS

### 4.1 *Berberis royleana* Extracted materials

The dried powder of crude methanol extracts obtained after filtration and rotary evaporation about 106 grams powder was found. Different solvents extracts were obtained after fractionation i.e. water fraction (55 g), *n*-hexane fraction (0.7 g), Ethyl acetate fraction (12 g) and chloroform (13 g) and 10 gm crude methanol extract was already stored.

### 4.2 Antibacterial activity of *Berberis royleana* Extract

Antibacterial activity of 100 μg *B. royleana* each (methanol, water, ethyl acetate, *n*-Hexane and chloroform) fraction extracts against 42 bacterial isolates showed decent activity against each bacterial isolates such bacterial isolates are *S. aureus* (16), *E. coli* (13*), K. pneumoniae* (3). *P. aeruginosa* (4), *Proteus spp* (3) and *S. Typhi* (3) which showed different zone of inhibitions (Table 4.2).

#### 4.2.1 Antibacterial activity of methanolic fraction

Hundred (100) μg of *B. royleana* methanolic extract showed maximum zone of inhibition (ZI) against *K. pneumoniae* i.e. 25.7 mm (SD ±1.5), while minimum ZI was found against *Proteus spp* 16.7 mm (SD ±3.21). The detailed description against all bacterial isolates is mentioned in Table 4.3. The results of methanolic extract against bacterial isolates have been compared with positive control (Ciprofloxacin 5 μg) as shown in figure 4.1. The DMSO was used as negative control.

#### 4.2.2 Antibacterial activity water fraction

Hundred (100) μg of *B. royleana* water fraction showed maximum zone of inhibition (ZI) against *K. pneumoniae* i.e. 24.4 mm (SD ±2.5), while minimum ZI was found against *Proteus spp* 15 mm (SD ±2.6). The detailed description against all bacterial isolates is mentioned in Table 4.4. The results of water extract against bacterial isolates have been compared with positive control (Ciprofloxacin 5 μg) as shown in figure 4.2. The DMSO was used as negative control.

#### 4.2.3 Antibacterial activity *n*-Hexane fraction

Hundred (100) μg of *B. royleana n*-hexane fraction showed maximum zone of inhibition (ZI) against *K. pneumoniae* i.e. 24.7 mm (SD ±1.5), while minimum ZI was found against *Proteus spp* 14 mm (SD ±3). The detailed description against all bacterial isolates is mentioned in Table 4.5. The results of *n*-hexane fraction against bacterial isolates have been compared with positive control (Ciprofloxacin 5 μg) as shown in figure 4.3. The DMSO was used as negative control.

#### 4.2.4 Antibacterial activity of Ethyl acetate fraction

Hundred (100) μg of *B. royleana* ethyl acetate fraction showed maximum zone of inhibition (ZI) against *E. coli* i.e. 16.6 mm (SD ±3.1), while minimum ZI was found against *Proteus spp* 12.7 mm (SD ±2.5). The detailed description against all bacterial isolates is mentioned in Table 4.6. The results of ethyl acetate fraction against bacterial isolates have been compared with positive control (Ciprofloxacin 5 μg) as shown in figure 4.4. The DMSO was used as negative control.

#### 4.2.5 Antibacterial activity of chloroform fraction

Hundred (100) μg of *B. royleana* chloroform fraction showed maximum zone of inhibition (ZI) against *S.* Typhi i.e. 21.6 mm (SD ±3.3), while minimum ZI was found against *Proteus spp* 13.3 mm (SD ±3.4). The detailed description against all bacterial isolates is mentioned in Table 4.7. The results of chloroform fraction against bacterial isolates have been compared with positive control (Ciprofloxacin 5 μg) as shown in figure 4.5. The DMSO was used as negative control.

## V. DISCUSSION

Microbial resistance is one of the major health issue in developing countries. Most of the bacterial and fungal species developed resistance to available drugs. An alternative remedy is essential to prevent this drug resistance as well as it should be cost effective. In current study examine the potential antimicrobial activity of methanolic, ethyl acetate, chloroform, n-hexane and water extracts of *B. royleana* against bacterial and fungal species by means of the whole plate and disc diffusion methods in order to compare the suitability of the identification methods and *in vitro* insecticidal activity of methanolic, ethyl acetate, chloroform, *n*-Hexane and water fraction extracts of *B. royleana* against insects by direct contact application and of all the activities of the extracts.

### 5.1 Antibacterial activity of methanolic fraction

In current research, 100 μg, methanolic fraction of *Berberis royleana* extract showed maximum ZI against *K. pneumoniae* as i.e. 25.7 mm (SD ±1.5) while that of minimum was against *Proteus spp.* i.e. 16.7 mm (SD ±3.21) followed by *S. Typhi* 25 mm (SD±1) and *S. aureus* 23 mm (SD ±2.7) *P. aeruginosa* 21.5 mm (SD ±5.1) and *E. coli* 20.62 mm (SD ±3.4). In similar report the crude methanolic extract *Berberis baluchistanica* showed antimicrobial activities against *Klebsiella pneumoniae* 12.67 (SD±0.76), *S. aureus* 17.33 (SD ±1.52), *E. coli* 12.33 (SD±0.58) *P. aeruginosa* 19.33 (SD±0.76) and *Salmonella* Typhi 10.67 (SD±1.04) [37]. Another research methanolic extract reveals ZI against *E. coli* (15 mm), *K. pneumoniae* (16 mm) and some pathogens are resistant i.e. *S. aureus* [38]. In another investigation methanolic root extracts of *Berberis lycium* exhibited maximum inhibitory zone *E. coli* (12 mm), *Pseudomonas spp* (11 mm), *Staphylococcus spp* (10 mm) [39]. In study Rasool *et al*., 2015 the antibacterial activity of methanol extract of *B. calliobotrys* against *S. aureus* (27±2.51) and *P. aeruginosa* (20±3) and results compared with ciprofloxacin [40]. Our studies are agreement with Kakar *et al.*, 2012 [37]. In current studies and the previous studies of (Rasool *et al*., 2015[38] [39] [40]) have good activity against bacterial species (gram negative and gram positive) with comparison of standard antibiotics but the some difference between both studies different number of bacterial isolates and used of different volume of concentrations.

### 5.2 Antibacterial activity of water fraction

In current research, 100 μg, water fraction of *Berberis royleana* extract showed maximum ZI i.e. 24.4 (SD ±2.5) against *K. pneumoniae* while minimum ZI was 15 (SD ±2.6) against *Proteus spp* followed by *S. Typhi* 23 (SD ±1) and *S. aureus* 21 (SD± 2.8), *E. coli* 18.9 (SD ±4) and *P. aeruginosa* 18.5 (SD ±3.8). In similar study the water fraction of *Berberis aristata* extract showed no activity against *Klebsiella pneumoniae, Proteus spp, P. aeruginosa, S.* Typhi, some other bacterial species but *S. aureus* (27.9±2.7) showed good activity [41]. In previous study of Malik *et al*., 2017 the aqueous extract showed antimicrobial activity against *Berberis artista* showed highest zone of inhibition for *E. coli* (10.0±1.3) followed by *S. typhimurium* (9.3±0.1) and *S. aureus* (8.3±1.7) [42]. In previous study of Dashti *et al.*, 2014 the aqueous extracts of the fruits of berberis showed antimicrobial activity against *S.* aureus ZI (11 ±0.87) and *E. coli* (10±0.87) [43]. Our results showed significant activity against all bacterial isolates as compare to previous studies [41] [42] [43] the reason may be isolation of the bacteria from different body sites and plant species. In present we found that previous studies have similar procedure but the plant species are different from the present study and different concentrations of extract are used against bacterial species and different standard drug.

### 5.3 Antibacterial activity of *n*- Hexane fraction

In current study, 100 μg, *n*-Hexane fraction of *Berberis royleana* extract showed maximum ZI i.e. 24.7 (SD ±1.5) against *K. pneumoniae* while that of minimum was against *Proteus spp.* i.e. 14 (SD ±3) followed by *S. Typhi* 24 (SD ±2), *P. aeruginosa* 22.5 (SD ±1) and *S. aureus* 20.1 (SD ±4.2). In similar study Shah *et al.*, 2012 analyzed for antibacterial activities, *n-*Hexane extract of *Berberis vulgaris* was exhibited no significant activity against the *Staphylococcus aureus, Klebsiella pneumonia* and *Escherichia coli* [4]. In Aziz *et al.*, 2011 study of *Vitex agnus-castus* hexane fraction demonstrated antibacterial activity against *B. cereus* (7.5 mm), *P. aeruginosa* (10mm) and *S. Typhi* (11mm) while other *Corynebacterium diptheriae*, *E. coli, K. pneumoniae, Proteus mirabills* and *S. aureus* show resistant [44]. Current research have significant activities against both Gram negative and gram positive bacterial isolates but the pervious [4] [44] studies there are some bacterial isolates were inactive to plant extract. There are same procedure and same fraction were used in previous studies as compared with present study.

### 5.4 Antibacterial activity Ethyl acetate fraction

In current study, 100 μg, ethyl acetate fraction of *Berberis royleana* extract showed maximum ZI i.e. 16.6 (SD ±3.1) against *E. coli* while that of minimum ZI i.e. 12.7 (SD ±2.5) against *Proteus spp* followed by *K. pneumoniae* 16.3 (SD ±1.5) and *S. aureus* 16.2 (SD ±2.8) and *P. aeruginosa* 14 (SD ±1.6). In similar study the ethyl acetate fraction extract of *Berberis baluchistanica* showed no activity against *K. pneumoniae, Proteus* and other bacterial species [45]. Another research reveals ZI against *E. coli* (15 mm), *K. pneumoniae* (16 mm) and some pathogens are resistant i.e. *S. aureus* [38]. Our results showed significant activity against all bacterial isolates as compare with [45], the reason may be isolation of the bacteria from different body sites and plant species. Our studies are agreement with Sasikumar *et al*. 2007 [38]. Both reports have similar procedure but the plant species are different from the present study and different concentrations (10, 5, 2.5 mg/ml) of ethyl acetate extract was used against bacterial species and different standard drug.

### 5.5 Antibacterial activity Chloroform fraction

In present investigation, 100 μg Chloroform fraction of *Berberis royleana* extract demonstrated maximum ZI i.e. 21.6 (SD ±3.3) against *S.* Typhi followed by *K. pneumoniae* 21.3 (SD ±3.3), *P. aeruginosa* 20.7 (SD ±3.4), *S. aureus* 19.7 (SD ±3.34), *E. coli* 18.5 (SD ±4.65) and minimum against *Proteus spp* 13.3 (SD ±3.51). In the [38] studies the chloroform root extracts of *B. tinctoria* exhibited maximum activity against *P. aeruginosa* (25 mm), *K. pneumoniae* (20 mm), *E. coli* (10 mm), *Salmonella* species (20 mm), *S. aureus* (15mm), *A. hydrophila* (15 mm) and other pathogens *V. cholera* and *V. parahemolyticus* were resistant. In study of [44] the chloroform fraction of *Vitex agnus-castus* exhibited ZI against *K. pneumoniae* (8 mm), *P. aeruginosa* (6.5mm) and *S. typhi* (10 mm) while *Corynebacterium diptheriae, E. coli, K. pneumoniae*, *P. mirabills* and *S. aureus* show resistant. In previous investigation of Baloch *et* al., 2013 Chloroform fraction of *Thuspeinanta brahuica* root showed strong activity against *B. subtilis* (28 mm) and moderate activity against *S. aureus* with (25 mm) of zone inhibition. *E. coli* showed (14 mm) of zone inhibition, *S.* Typhi showed (12 mm) of zone inhibition whereas *P. aeruginosa* was resistant to chloroform fraction [46]. Our studies are in agreement with [38]. In present study significant activities against all bacterial isolates were found, as compared to previous studies [44] [46] where some extracts showed activities, however, some of them were inefficient against resistant bacterial species. The reason may be geographical distribution, site of sample collection.

## VI. CONCLUSIONS AND RECOMMENDATIONS

### CONCLUSIONS

Antimicrobial activities of *Berberis royleana* methanol and water fractions showed promising results as compared to chloroform, ethyl acetate and n-hexane the maximum ZI against *S. aureus, E. coli, S.* Typhi, *P. aeruginosa, K. pneumoniae* recorded was 25.7 ±1.5 mm while these extract showed minimum ZI against *Proteus spp* from 16.7 ±3.21 mm.

From current report it may be concluded that *B. royleana* extracts have medicinally effective potentials against drugs resistant bacteria.

### RECOMMENDATIONS

1. Fourier Transform Infrared Radiation (FTIR) and Gas Chromatographic Mass Spectroscopy (GCMS) analysis are highly recommended to explore the active chemicals in *B. royleana.*
2. The nanoparticles may be prepared using *B. royleana* to test against MDR bacteria.

## VIII. ANNEXER

**Table.4.1.** Various Extracts of *Berberis royleana*.

| S. No. | Extracts / Fractions | Extracted powder in grams |
| --- | --- | --- |
| 1 | Crude methanol | 106 g |
| 2 | Water | 55 g |
| 3 | *n*- Hexane | 0.7 g |
| 4 | Ethyl acetate | 12 g |
| 5 | Chloroform | 13 g |

**Table.4.2.** Frequency distribution of bacterial isolates.

| S. No | Bacterial isolates | Number of isolates |
| --- | --- | --- |
| 1 | *Staphylococcus aureus* | 16 |
| 2 | *Escherichia coli* | 13 |
| 3 | *Pseudomonas aeruginosa* | 4 |
| 4 | *Klebsiella pneumoniae* | 3 |
| 5 | *Proteus spp* | 3 |

**Figure.4.1.**
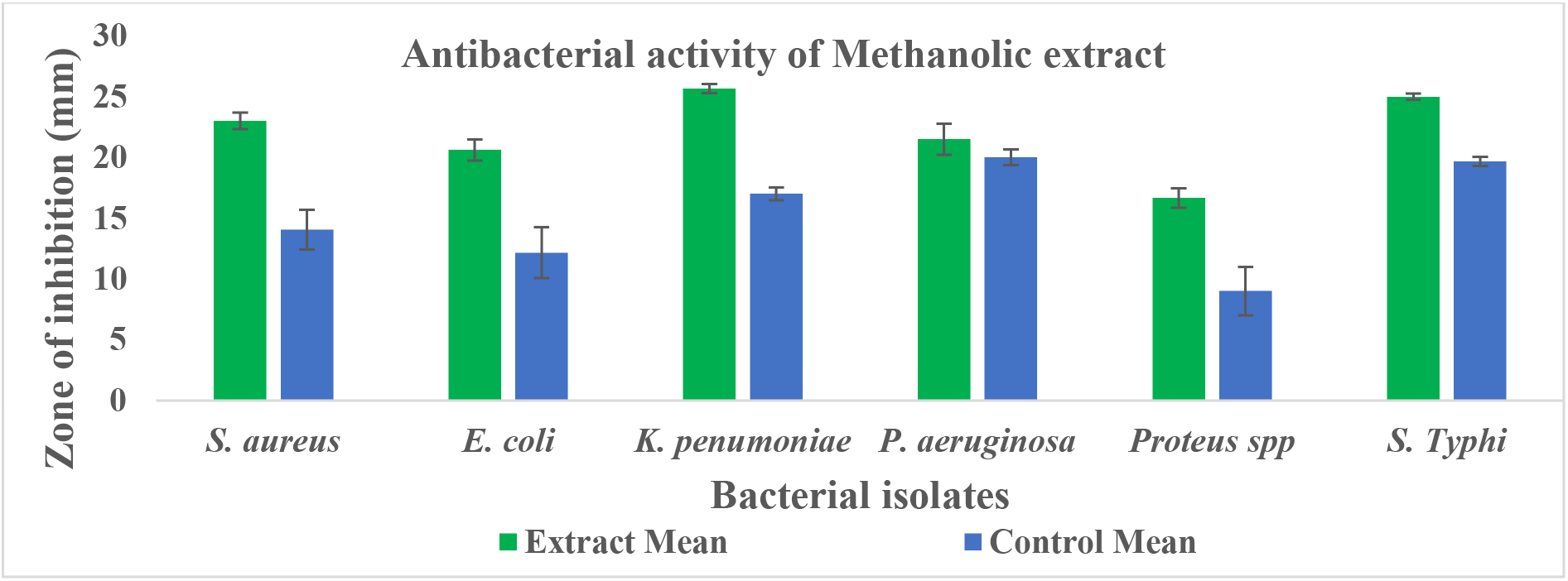
Methanol fraction antibacterial activity *B. royleana* extract. Methanolic extract 100 μg and positive control ciprofloxacin 5 μg

**Table.4.3.** Methanolic fraction and antibiotic antibacterial activity.

| S. No. | Bacterial isolate | Extract Mean ZI ± SD | Control Mean ZI ± SD |
| --- | --- | --- | --- |
| 1 | *S. aureus* | 23 ±2.7 | 14.1 ±6.56 |
| 2 | *E. coli* | 20.62 ±3.4 | 12.2 ±8.36 |
| 3 | *K. pneumoniae* | 25.7 ±1.5 | 17 ±1 |
| 4 | *P. aeruginosa* | 21.5 ±5.1 | 20 ±2.58 |
| 5 | *Proteus spp* | 16.7 ±3.21 | 9 ±7.9 |
| 6 | *S. Typhi* | 25 ±1 | 19.7 ±1.53 |

**Figure.4.2.**
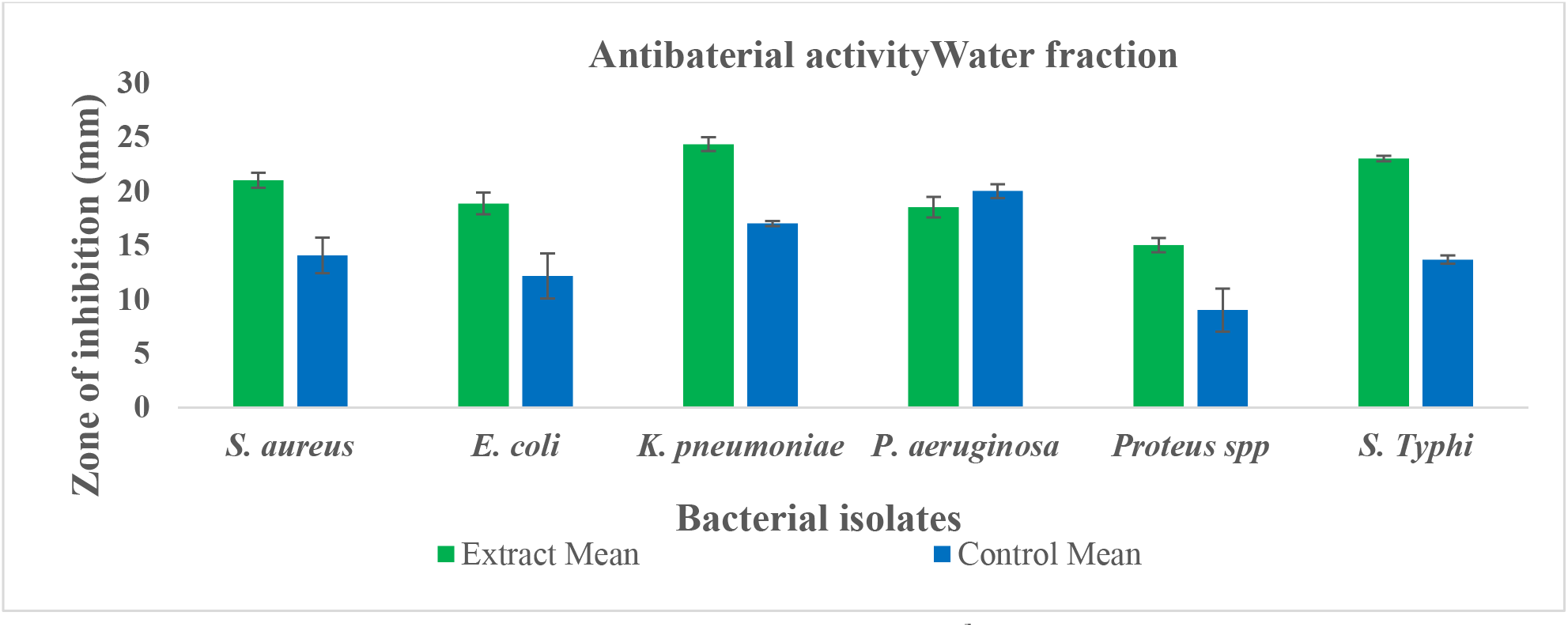
Water fraction antibacterial activity of *B. royleana* extract. Water extract 100 μg and positive control ciprofloxacin 5 μg

**Table.4.4.**
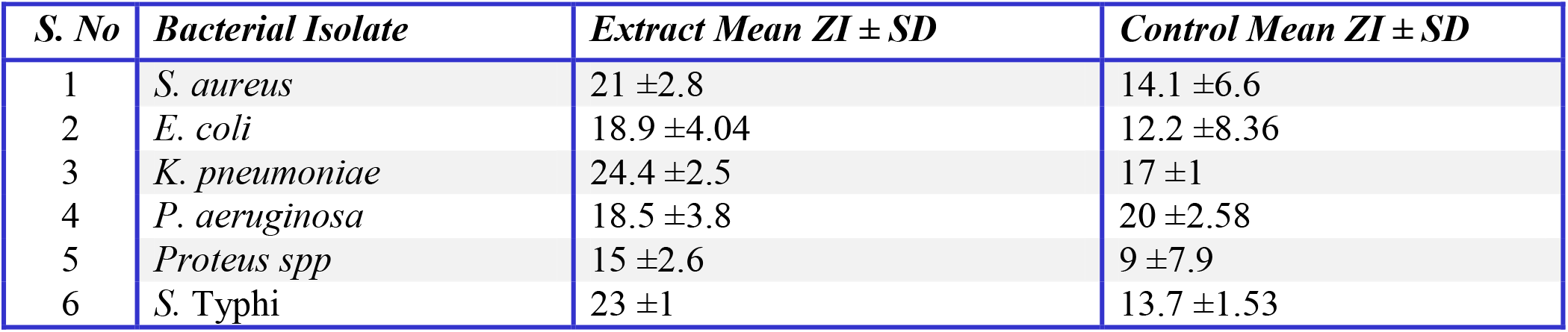
Water fraction and antibiotic antibacterial activity.

| *S. No* | *Bacterial Isolate* | *Extract Mean ZI ± SD* | *Control Mean ZI ± SD* |
| --- | --- | --- | --- |
| 1 | *S. aureus* | 21 ±2.8 | 14.1 ±6.6 |
| 2 | *E. coli* | 18.9 ±4.04 | 12.2 ±8.36 |
| 3 | *K. pneumoniae* | 24.4 ±2.5 | 17 ±1 |
| 4 | *P. aeruginosa* | 18.5 ±3.8 | 20 ±2.58 |
| 5 | *Proteus spp* | 15 ±2.6 | 9 ±7.9 |
| 6 | *S. Typhi* | 23 ±1 | 13.7 ±1.53 |

**Figure.4.3.**
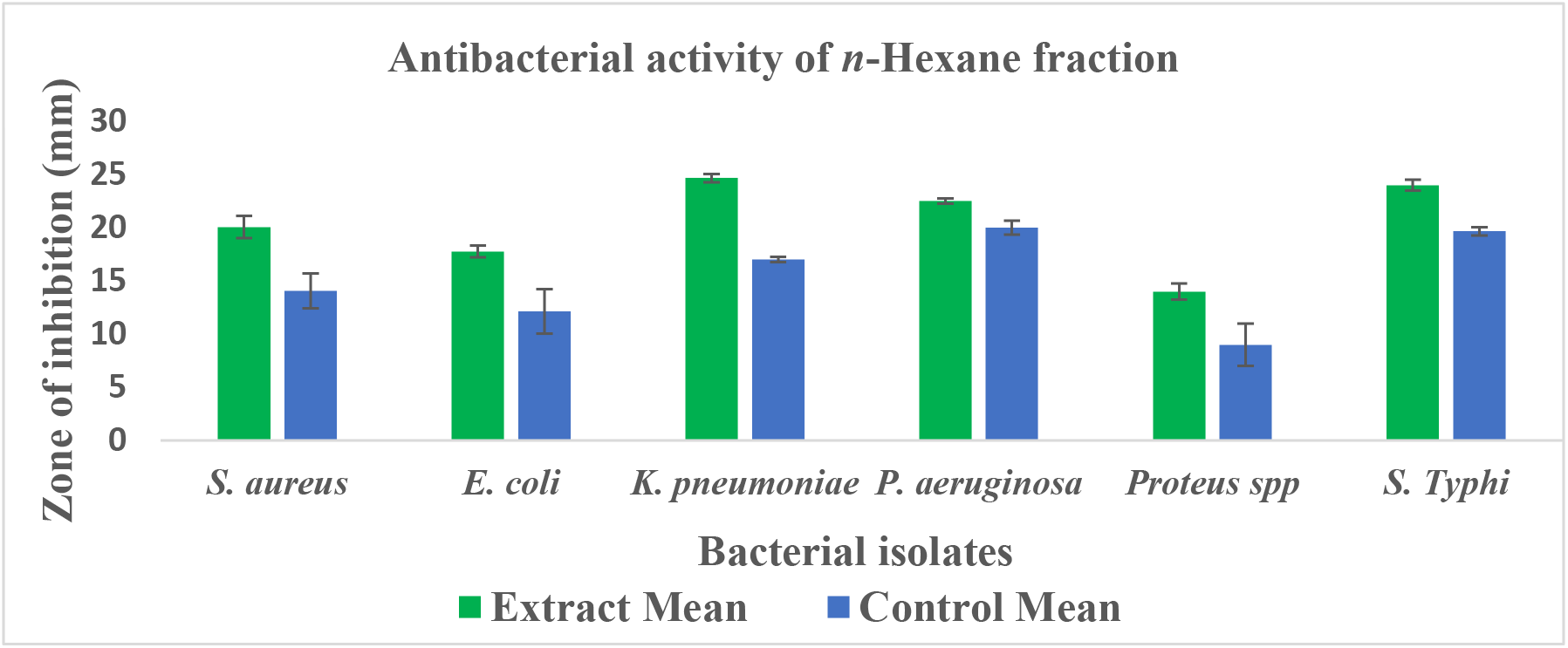
*n*- Hexane fraction antibacterial activity of *B. royleana* extract. *n*-Hexane extract 100 μg and positive control ciprofloxacin 5 μg

**Table No.4.5.** *n*- Hexane fraction and antibiotic antibacterial activity.

| *S. No* | *Bacterial isolate* | *Extract Mean ZI ± SD* | *Control Mean ZI ± SD* |
| --- | --- | --- | --- |
| 1 | *S. aureus* | 20.1 ±4.2 | 14.1 ±6.5 |
| 2 | *E. coli* | 17.8 ±2.16 | 12.2 ±8.36 |
| 3 | *K. pneumoniae* | 24.7 ±1.5 | 17 ±1 |
| 4 | *P. aeruginosa* | 22.5 ±1 | 20 ±2.58 |
| 5 | *Proteus spp* | 14 ±3 | 9 ±7.9 |
| 6 | *S. Typhi* | 24 ±2 | 19.7 ±1.52 |

**Figure.4.4.**
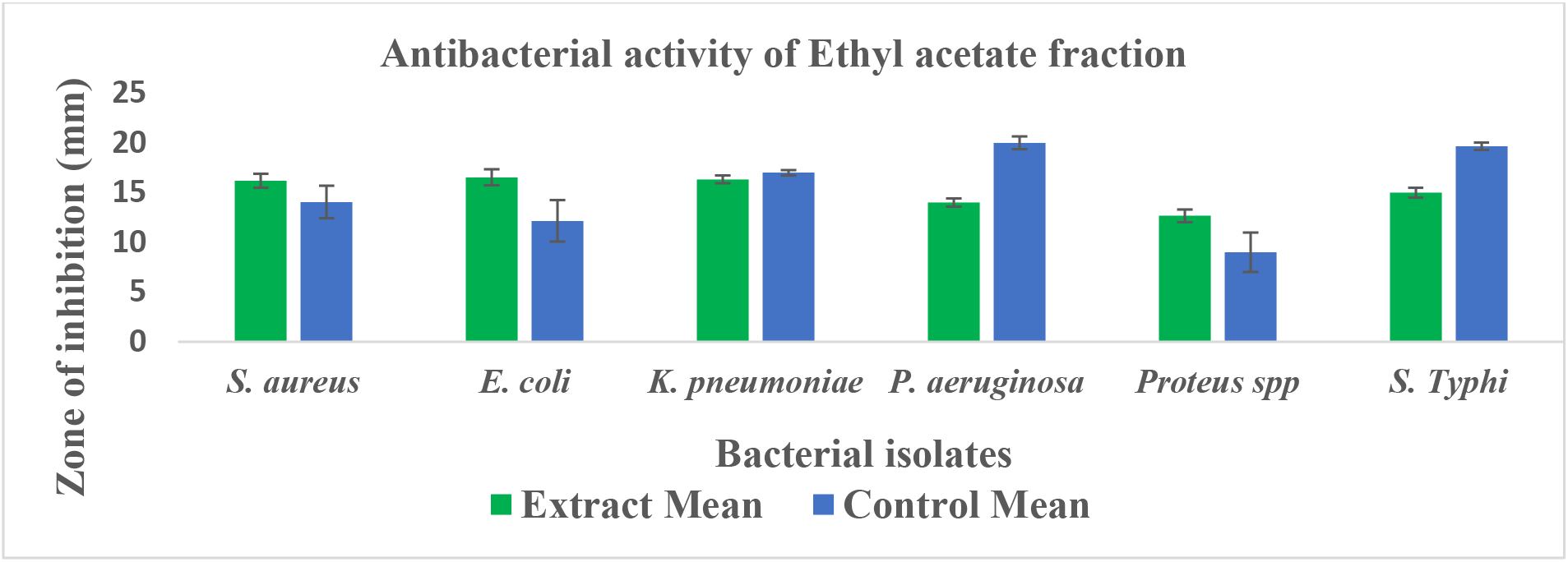
Ethyl acetate fraction antibacterial activity of *B. royleana* extract. Ethyl acetate extract 100 μg and positive control ciprofloxacin 5 μg

**Table.4.6.** Ethyl acetate fraction and antibiotic antibacterial activity.

| *S. No* | *Bacterial Isolate* | *Extract Mean ZI ± SD* | *Control Mean ZI ± SD* |
| --- | --- | --- | --- |
| 1 | *S. aureus* | 16.2 ± 2.85 | 14.2 ± 6.56 |
| 2 | *E. coli* | 16.6 ± 3.2 | 12.2 ± 8.36 |
| 3 | *K. pneumoniae* | 16.3 ± 1.53 | 17 ± 1 |
| 4 | *P. aeruginosa* | 14.0 ± 1.63 | 20 ± 2.6 |
| 5 | *Proteus spp* | 12.7 ± 2.52 | 9 ± 7.93 |
| 6 | *S. Typhi* | 15.0 ± 2 | 19.7 ± 1.53 |

**Figure.4.5.**
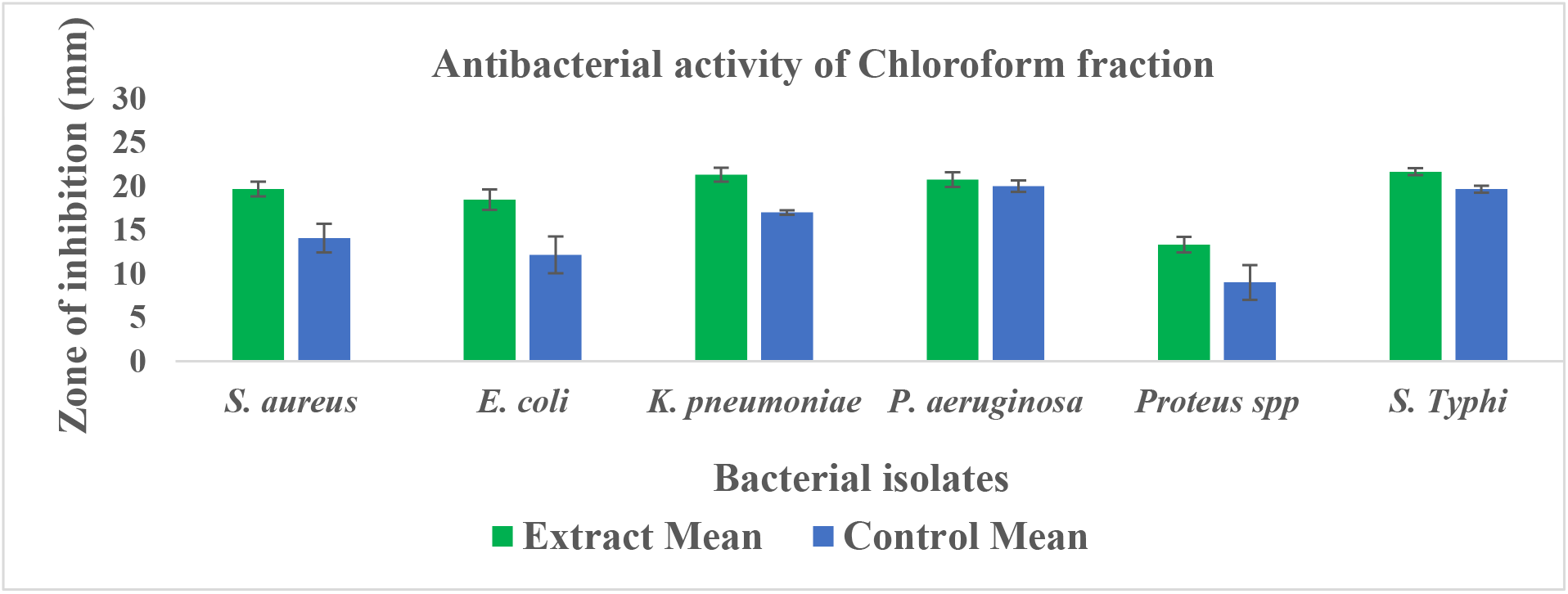
Chloroform fraction antibacterial activity of *B. royleana*. Chloroform extract 100 μg and positive control ciprofloxacin 5 μg

**Table. 4.7.**
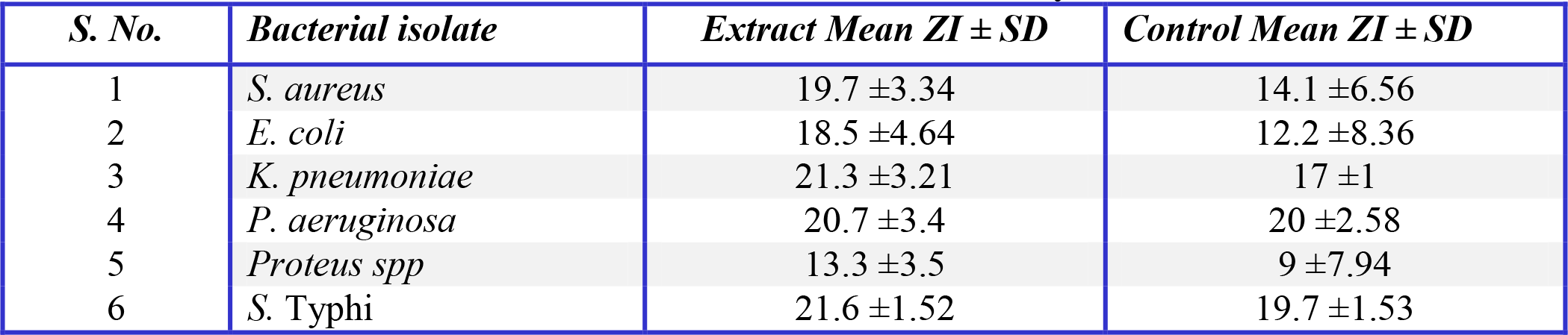
Chloroform fraction and antibiotic antibacterial activity.

| *S. No.* | *Bacterial isolate* | *Extract Mean ZI ± SD* | *Control Mean ZI ± SD* |
| --- | --- | --- | --- |
| 1 | *S. aureus* | 19.7 ±3.34 | 14.1 ±6.56 |
| 2 | *E. coli* | 18.5 ±4.64 | 12.2 ±8.36 |
| 3 | *K. pneumoniae* | 21.3 ±3.21 | 17 ±1 |
| 4 | *P. aeruginosa* | 20.7 ±3.4 | 20 ±2.58 |
| 5 | *Proteus spp* | 13.3 ±3.5 | 9 ±7.94 |
| 6 | *S. Typhi* | 21.6 ±1.52 | 19.7 ±1.53 |

**Figure.4.9.**
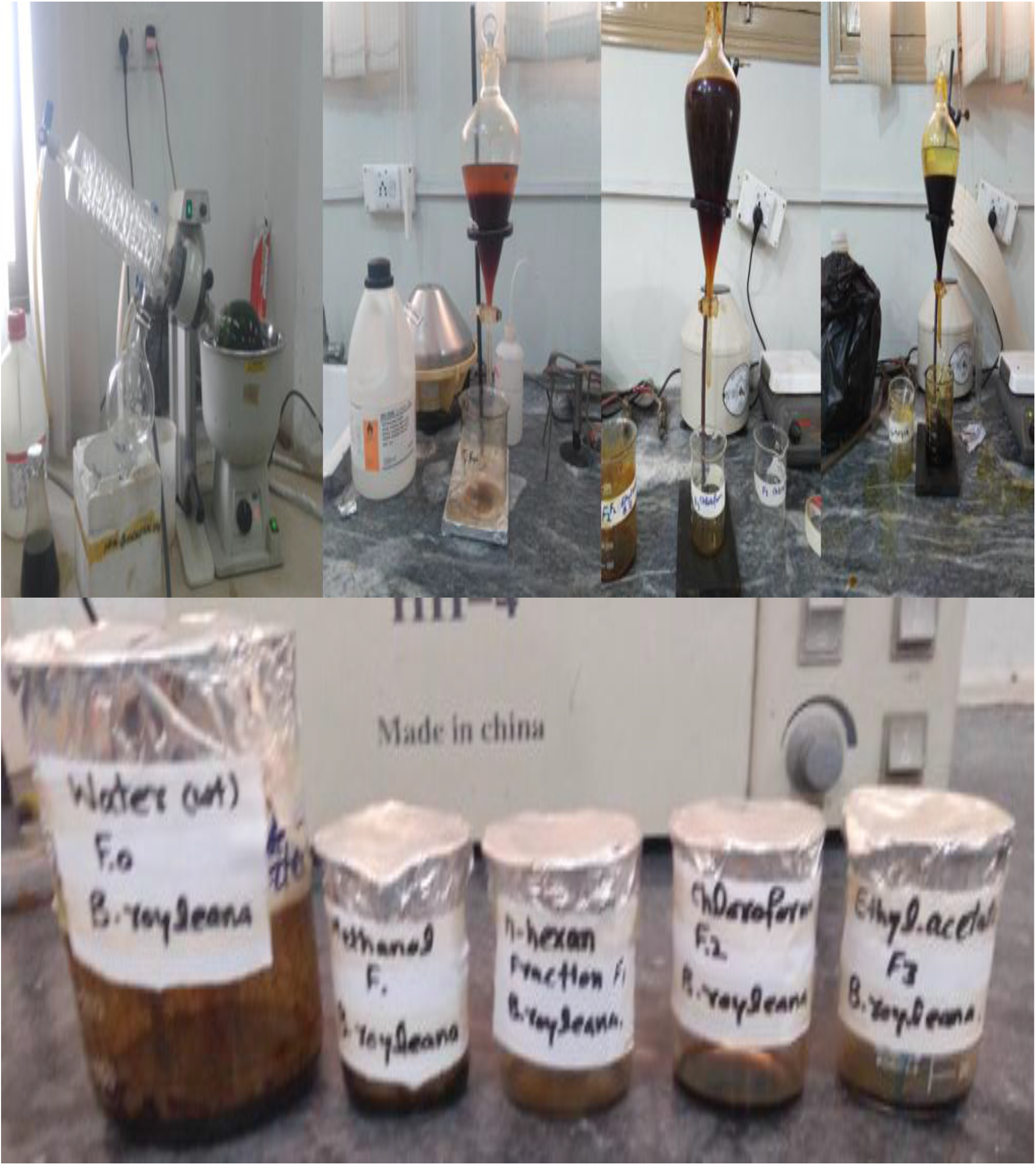
Extracts and their fractions of *Berberis royleana*.

**Figure.4.10.**
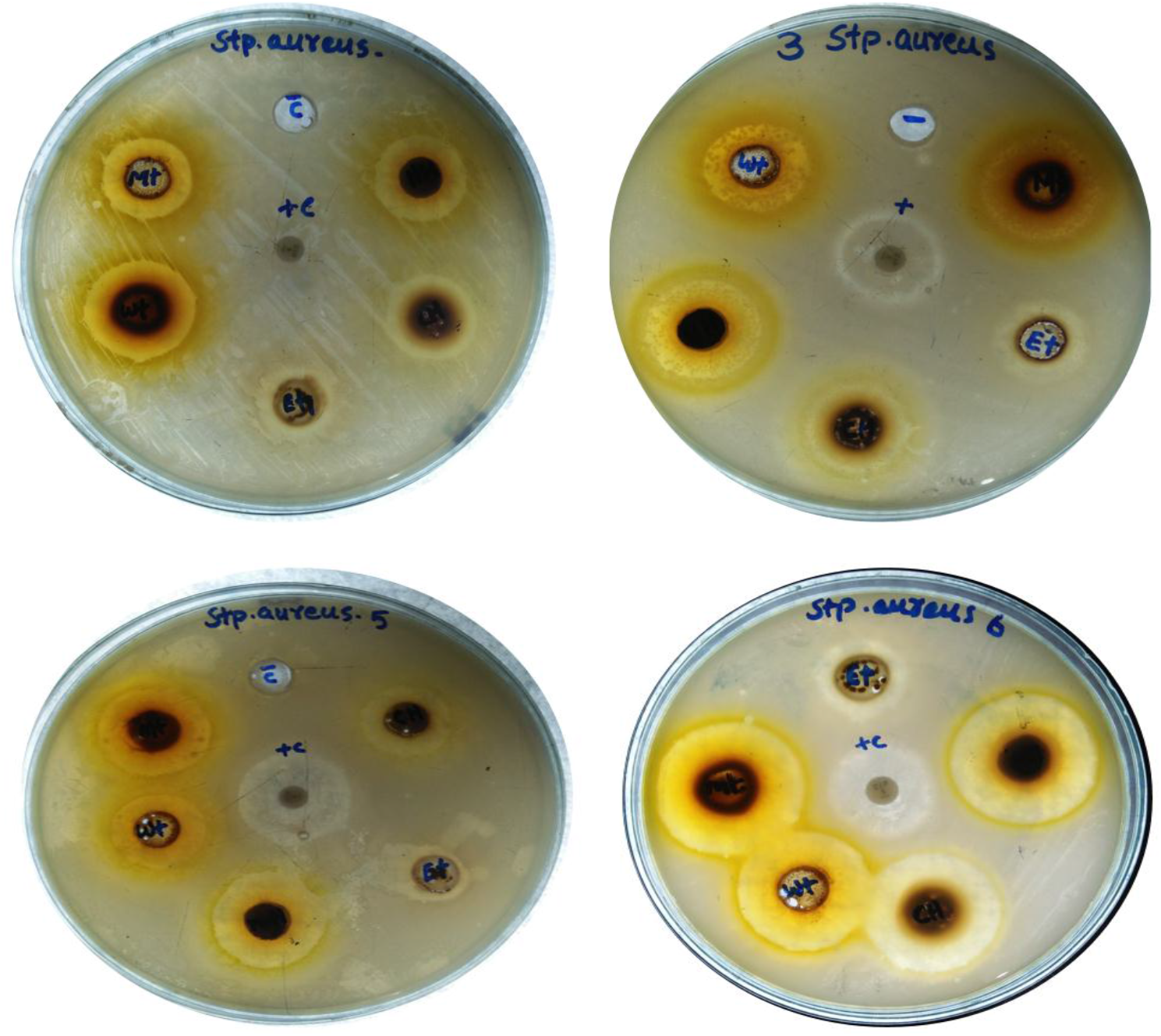
Antibacterial activity of *B. royleana* extract against *Staphylococcus aureus*.

**Figure.4.11.**
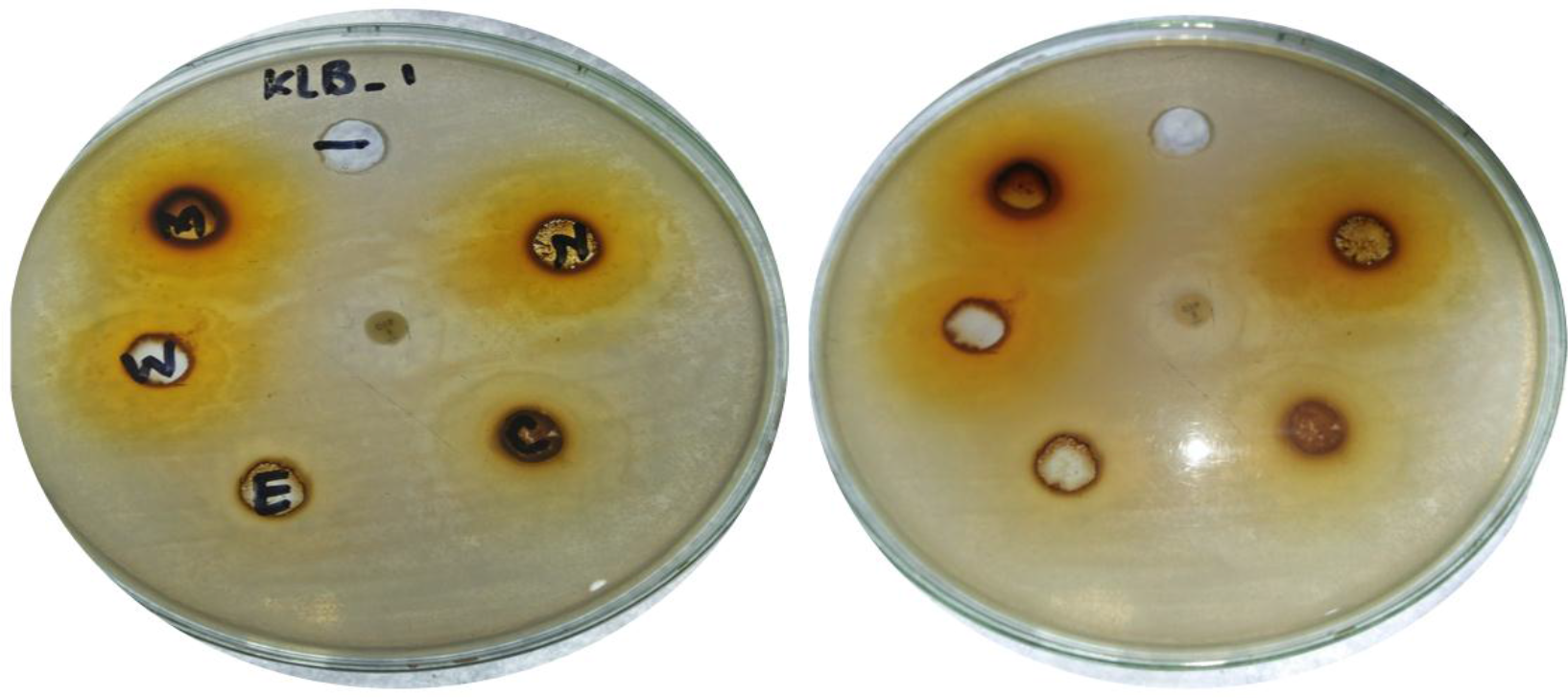
Antibacterial activity of *B. royleana* extract against *Klebsiella pneumoniae*.

**Figure.4.12.**
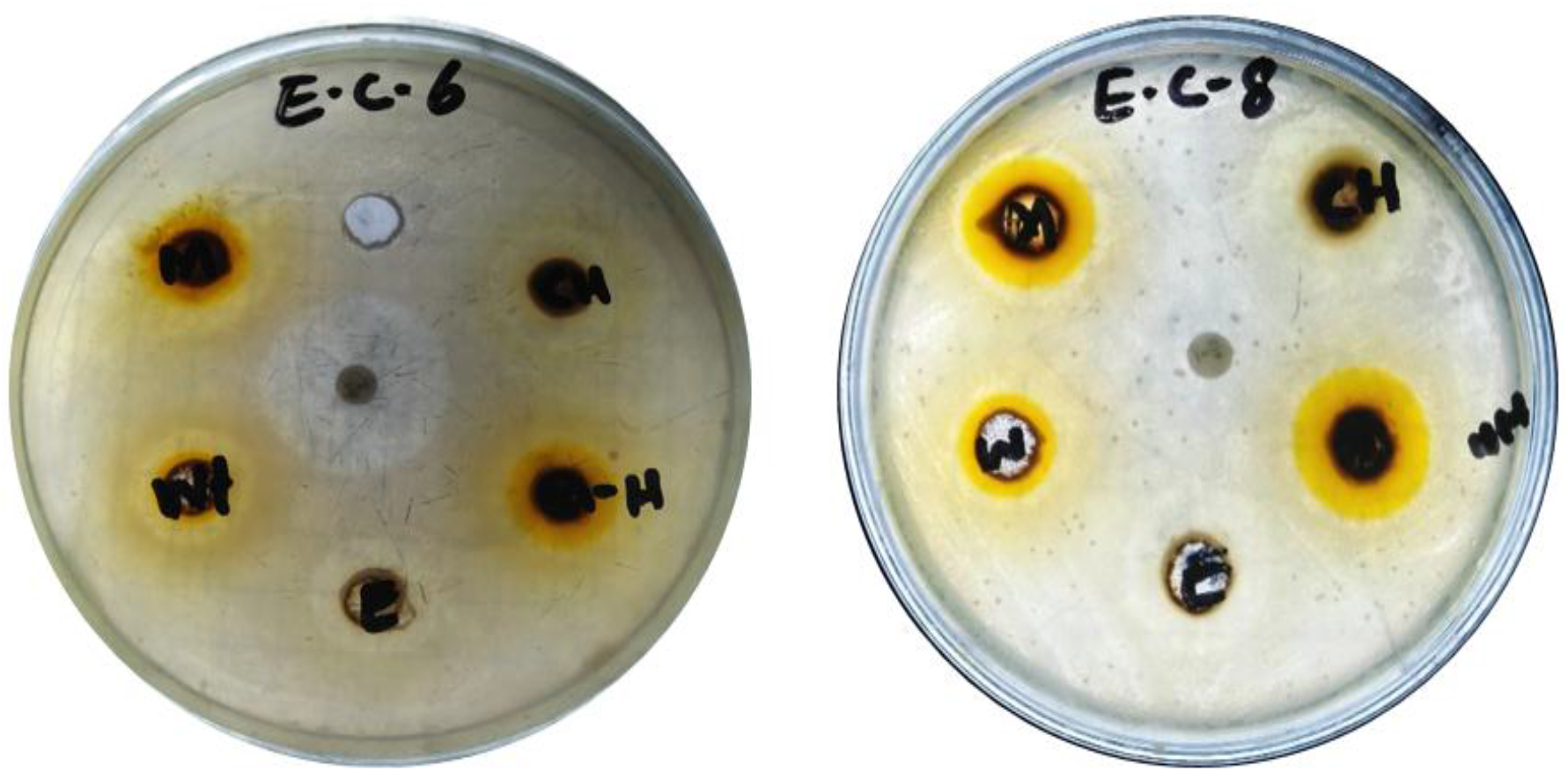
Antibacterial activity of *B. royleana* extract against *Escherichia coli*.

**Figure.4.13.**
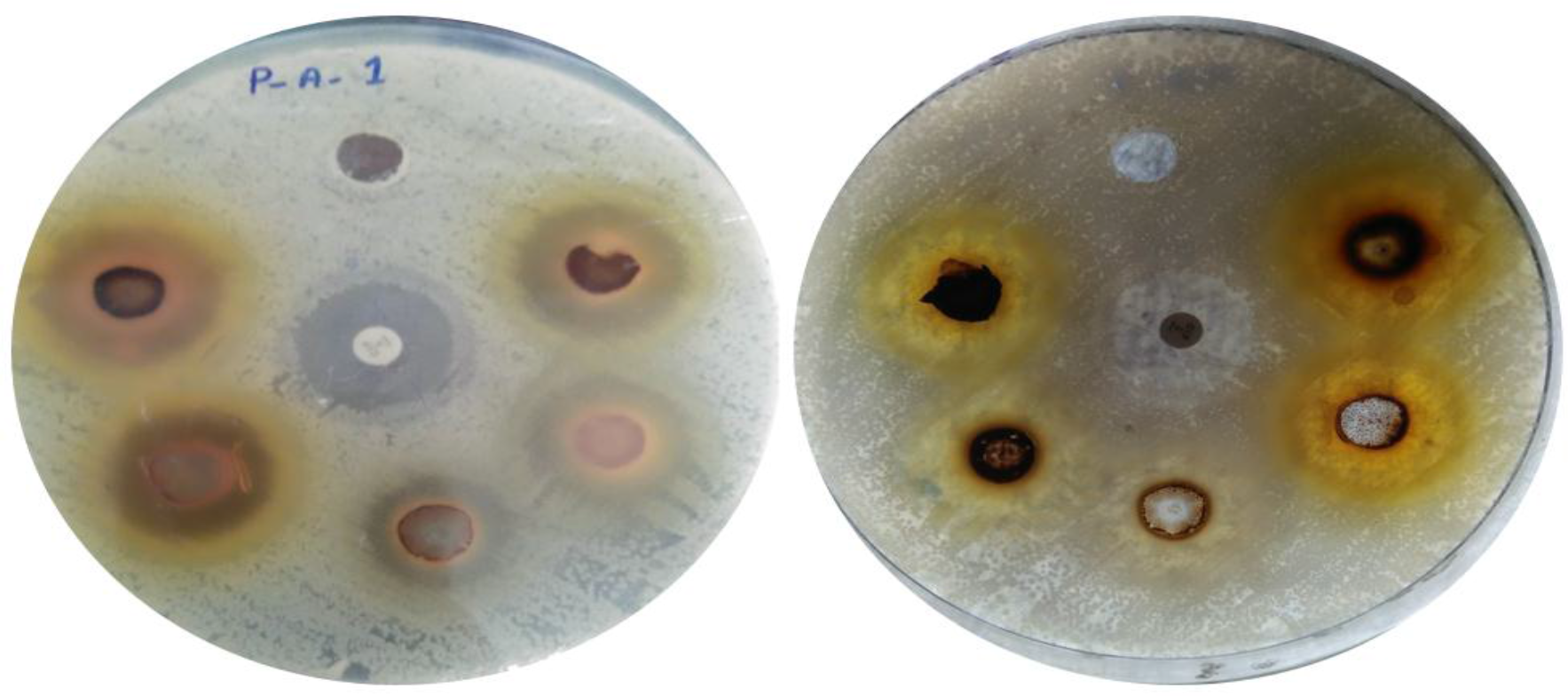
antibacterial activity of *B. royleana* extract against *Pseudomonas aeruginosa*.

**Figure.4.14.**
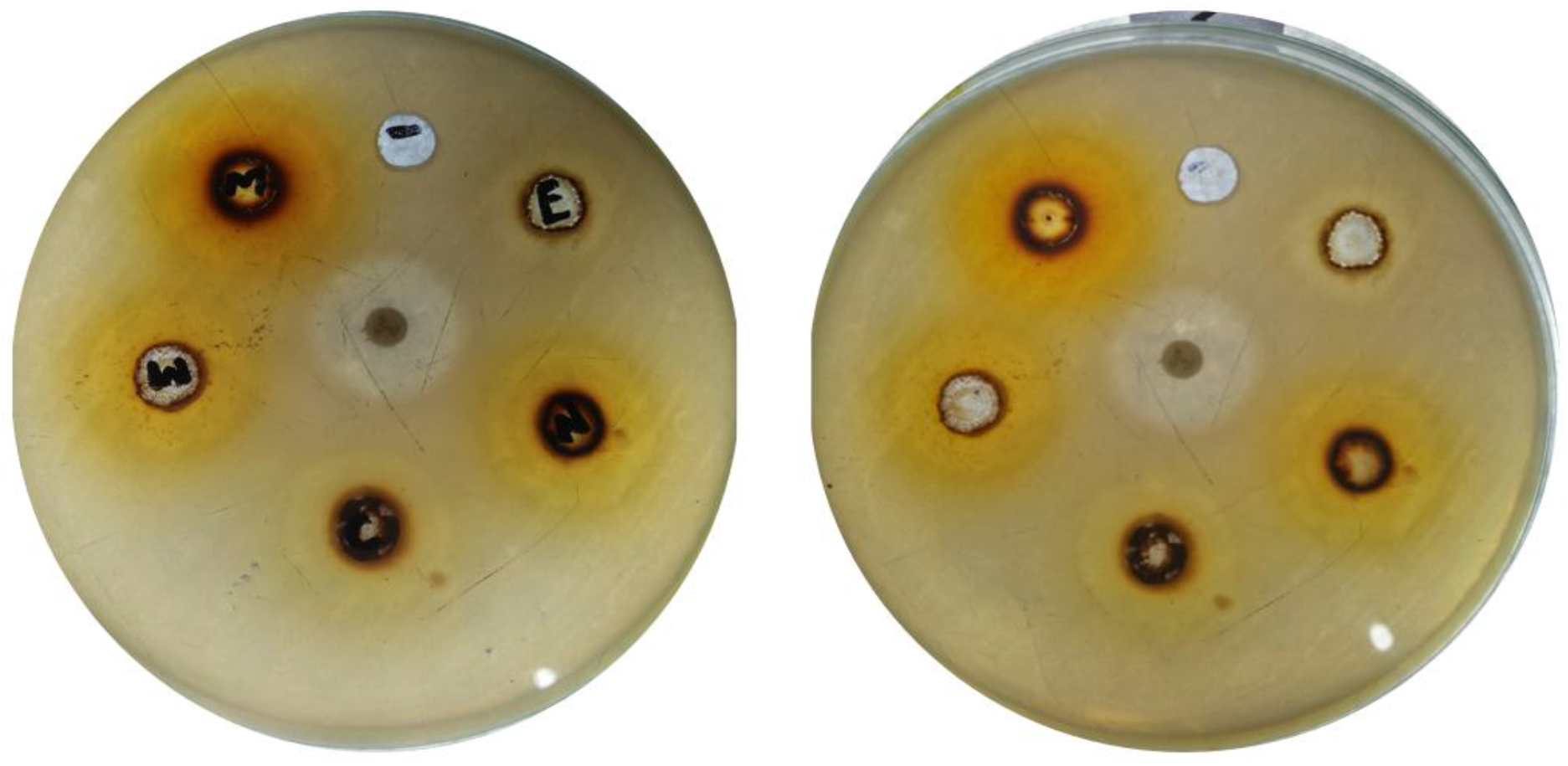
Antibacterial activity of *B. royleana* extract against *Salmonella* Typhi.

**Figure.4.15.**
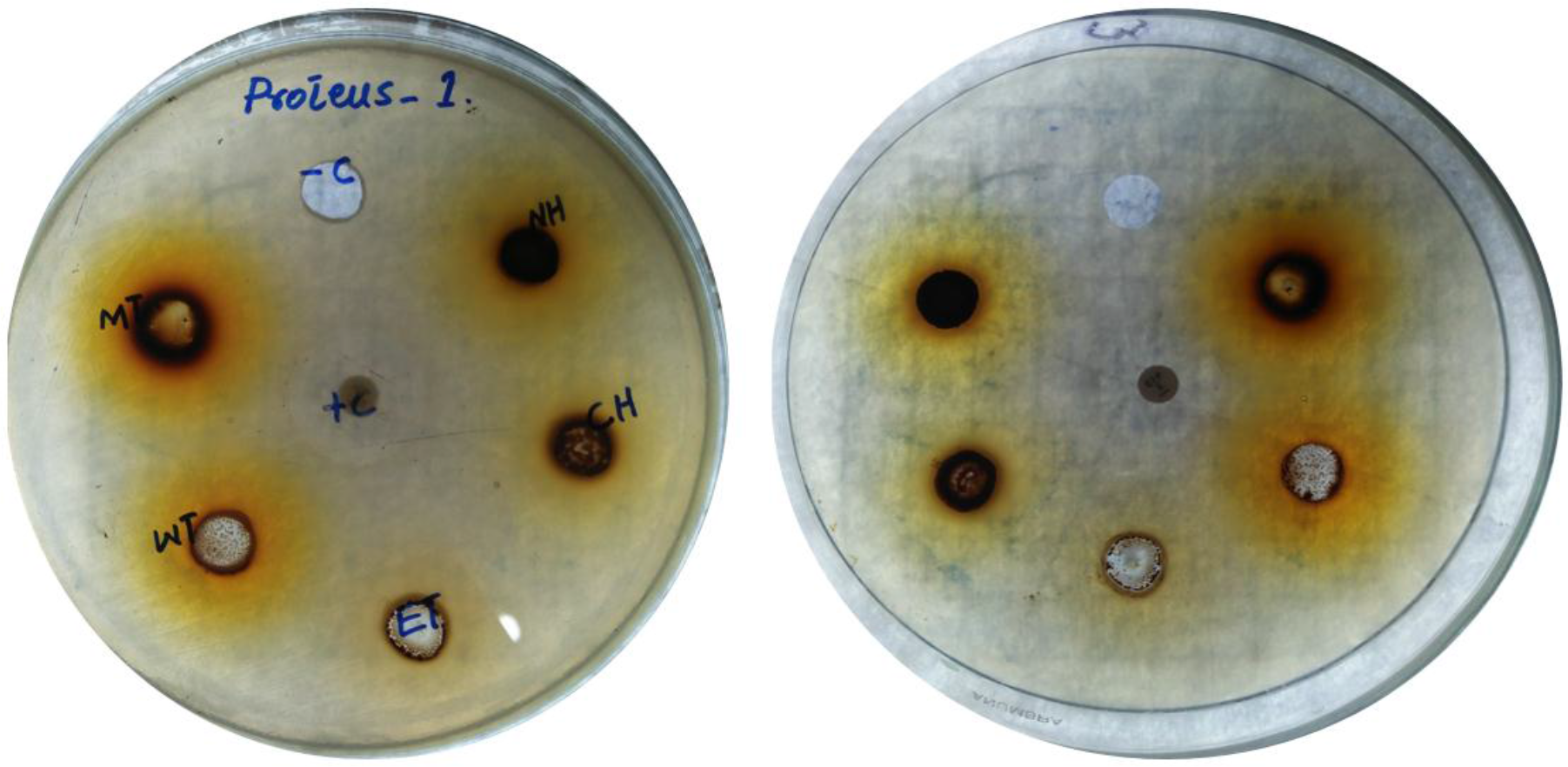
Antibacterial activity of *Berberis royleana* extract against *Proteus spp.*

